# mGluR5 Negative Modulators for Fragile X: Resistance and Persistence

**DOI:** 10.1101/2021.07.02.450894

**Authors:** David C Stoppel, Patrick K McCamphill, Rebecca K Senter, Arnold J Heynen, Mark F Bear

## Abstract

Fragile X syndrome (FXS) is caused by silencing of the human *FMR1* gene and is the leading monogenic cause of intellectual disability and autism. Abundant preclinical data indicated that negative allosteric modulators (NAMs) of metabotropic glutamate receptor 5 (mGluR5) might be efficacious in treating FXS in humans. Initial attempts to translate these findings in clinical trials have failed, but these failures provide the opportunity for new discoveries that will improve future trials. The emergence of acquired treatment resistance (“tolerance”) after chronic administration of mGluR5 NAMs is a potential factor in the lack of success. Here we confirm that FXS model mice display acquired treatment resistance after chronic treatment with the mGluR5 NAM CTEP in three assays commonly examined in the mouse model of FXS: (1) audiogenic seizure susceptibility, (2) sensory cortex hyperexcitability, and (3) hippocampal protein synthesis. Cross-tolerance experiments suggest that the mechanism of treatment resistance likely occurs at signaling nodes downstream of glycogen synthase kinase 3α (GSK3α), but upstream of protein synthesis. The rapid emergence of tolerance to CTEP begs the question of how previous studies showed an improvement in inhibitory avoidance (IA) cognitive performance after chronic treatment. We show here that this observation was likely explained by timely inhibition of mGluR5 during a critical period, as brief CTEP treatment in juvenile mice is sufficient to provide a persistent improvement of IA behavior measured many weeks later. These data will be important to consider when designing future fragile X clinical trials using compounds that target the mGluR5-to-protein synthesis signaling cascade.

## Introduction

Fragile X syndrome (FXS) is the most common monogenic intellectual disability disorder, affecting approximately 1:4000 males and 1:8000 females [1]. 60−75% of boys and 20−40% of girls with FXS are diagnosed with autism spectrum disorder (ASD), making FXS the most prevalent monogenic cause of ASD [2]. In almost all cases, FXS arises from a CGG trinucleotide repeat expansion in the 5` untranslated region of the fragile X mental retardation gene (*FMR1*) which fully silences expression of its protein product, fragile X mental retardation protein (FMRP) [3]. Decades of preclinical research have identified myriad disruptions to brain function in genetically engineered animal models of FXS, greatly advancing our understanding of the underlying molecular and cellular disease mechanisms [4].

Preclinical research has identified two broad classes of pathophysiological mechanisms related to two well described neuronal functions of FMRP: (1) altered protein synthesis regulation caused by loss of FMRP binding to mRNA [5; 6; 7; 8; 9; 10; 11; 12; 13], and (2) altered ion channel function caused by the disruption of FMRP protein-protein interactions with ion channels [9; 14; 15; 16; 17; 18; 19]. These two disease mechanisms impair synapse development, alter the balance between synaptic excitation and inhibition, and cause widespread increases in neuronal activity [20]. To make matters more complicated, the hyperexcitability caused by these proximal molecular defects can feed back to further alter protein synthesis, ion channel function, and synapse development. Short of gene therapy, strategies to tamp down neuronal hyperexcitability and altered proteostasis offer the best prospects to improve the course of FXS, particularly if initiated early in development.

The sheer diversity of ion channels involved in FXS pathology likely limits the potential benefit of targeting any single channel type [21]. On the other hand, the myriad consequences of altered protein synthesis regulation in FXS suggested that targeting this process has the potential to confer broad phenotypic improvement. This line of reasoning provided the rationale behind the “mGluR theory of fragile X”, as metabotropic glutamate receptor 5 (mGluR5) is widely expressed in the forebrain and known to modulate neuronal protein synthesis [22; 23]. Indeed, extensive studies in multiple animal models of FXS have shown that diverse disease phenotypes are corrected by inhibiting mGluR5 or key signaling nodes downstream of this receptor [8; 24; 25]. In addition to altered protein synthesis, these phenotypes include but are not restricted to hyperexcitability in sensory neocortex, increased cortical dendritic spine density, epileptiform activity in the hippocampus, audiogenic seizures (AGS), and impaired cognition measured by performance in an inhibitory avoidance (IA) task.

These encouraging animal studies led to human clinical trials using negative allosteric modulators (NAMs) of mGluR5 (mavoglurant and basimglurant). However, these studies failed to demonstrate improvements in the primary therapeutic endpoints [26]. The reasons for these failures have been extensively discussed and may include inadequate measures of target engagement and treatment response, suboptimal selection of drug doses and treatment duration, and the older age of the subjects [8; 27; 28]. There is also anecdotal evidence that although benefits were observed initially, the effectiveness of treatment faded with chronic dosing (https://www.fraxa.org/fragile-x-clinical-trials-mglur-theory/).

Resolving the obvious disconnect between the robust and highly reproducible rescue achieved in fragile X animal models and the failure to observe efficacy in humans is of enormous importance. Here we focus on one potential explanation for this discrepancy; the development of acquired treatment resistance (“tolerance”) that emerges during chronic drug treatment, a common obstacle for neuropsychiatric drug development [29]. Our study builds on previous observations of diminished mGluR5 NAM effectiveness with repeat dosing in the audiogenic seizure assay [30]. In that early work, the protection from AGS conferred by a single *in vivo* injection of the mGluR5 NAMs MPEP was significantly reduced (but not eliminated) after repeated daily dosing. This effect appears to be exacerbated when MPEP is combined with a GABA-B receptor agonist [31] and after dosing with CTEP, another mGluR5 NAM with greater selectivity and a much longer half-life [32]. In the current study we set out to address three key questions that emerge from these findings. (1) Is acquired treatment resistance to chronic mGluR5 NAM treatment observed in other FXS phenotypic assays? (2) Does resistance to treatment with an mGluR5 NAM impact the effectiveness of an intervention further downstream in the signaling pathway? (3) What accounts for the observation that in assays such as inhibitory avoidance, months long treatment with mGluR5 NAMs corrected behavior?

Our data show acquired treatment resistance following chronic treatment with the mGluR5 NAM CTEP occurs not only in AGS but also in assays of visual cortical hyperexcitability and elevated basal hippocampal protein synthesis. Treatment resistance is not due to increased mGluR5 expression or sensitivity as it cannot be overcome by additional treatment with structurally distinct mGluR5 NAMs. Rather, the mechanism of acquired treatment resistance appears to be downstream of the Ras-ERK1/2 pathway that links mGluR5 activation with increased synaptic protein synthesis. A selective inhibitor of GSK3α was unable to overcome acquired treatment resistance, but a protein synthesis inhibitor was still effective, positioning the mechanism of acquired treatment resistance between GSK3α and translation initiation. Finally, our data reveal that correction of IA deficits in adult mice does not require chronic dosing at all, but rather can be achieved by brief but timely treatment earlier in postnatal development.

Taken together these results provide additional evidence that acquired treatment resistance occurs following chronic mGluR5 inhibition and broadly impacts the ability of these drugs to correct pathophysiological phenotypes. However, treatment resistance is not an impediment to phenotypic improvements if the pharmacological interventions are timed to occur during critical developmental windows. These findings will be important to consider during study design for future clinical trials in FXS testing compounds that target mGluR5 and the downstream signaling cascade linked to altered proteostasis.

## Materials and Methods

### Study Design

Male *Fmr1* knockout (Fmr1-KO) and wildtype (WT) littermates on the C57BL/6J background were studied in all experiments. The breeding scheme was female Fmr1 heterozygous mice (Jackson Laboratory Stock Number: 003025) crossed with wildtype male C57BL/6J mice (Jackson Laboratory Stock Number: 000664). Sample size was determined by a power analysis or laboratory historical experience and no outliers were removed from any data sets. Age-matched littermates were randomized to treatment groups and a balanced number of Fmr1-KO and WT mice were used. Mice were group housed on static racks and maintained on a 12:12 hour light:dark cycle. The Committee on Animal Care at MIT approved all experimental techniques, and all animals were handled in accordance with NIH and MIT guidelines.

### Reagents

The mGluR5 specific negative allosteric modulator CTEP (chloro-4-((2,5-dimethyl-1-(4-(trifluoromethoxy)phenyl)-1H-imidazol-4-yl)ethynyl)pyridine) was purchased from Sigma-Aldrich (SML2306) and administered at 2 mg/kg concentration by intraperitoneal (i.p.) injection. CTEP was prepared daily as a microsuspension in vehicle (0.9% NaCl, 0.3% Tween-80). The GSK3α selective inhibitor BRD0705 was synthesized at the Broad Institute at MIT and confirmed to be of ≥ 95% purity based on HPLC LC-MS and 1H NMR analysis and administered at 30 mg/kg by i.p. injection. BRD0705 was prepared daily from frozen 50 mM stocks in DMSO. The mGluR5 specific negative allosteric modulator MPEP hydrochloride (2-Methyl-6-(phenylethynyl)pyridine hydrochloride) was purchased from Tocris (1212) and administered to brain slices at 30 μM by bath perfusion. MPEP was prepared daily in vehicle (100% aCSF, see below). The mGluR5 specific negative allosteric modulator MTEP (3-((2-Methyl-1,3-thiazol-4-yl)ethynyl)pyridine hydrochloride) was purchased from Tocris (2921) and administered to brain slices at 1 μM by bath perfusion. MTEP was prepared daily in vehicle (100% aCSF, see below). The translation elongation inhibitor cycloheximide (3-[2-(3,5-Dimethyl-2-oxocyclohexyl)-2-hydroxyethyl]glutarimide; CHX) was purchased from Sigma-Aldrich (C7698) and administered at 120 mg/kg concentration by i.p. injection and to brain slices at 60 μM by bath perfusion. For i.p. injection CHX was prepared daily in vehicle (0.9% NaCl). For bath application CHX was prepared daily in vehicle (aCSF, see below).

### Audiogenic seizure assay

AGS experiments were performed as previously described [32]. Fmr1-KO and WT mice were housed on static racks to prevent auditory desensitization that occurs with chronic exposure to the ambient noise of ventilated racks. For acute dosing experiments, mice received i.p. injections with vehicle or drug in a separate room 1-2 hours prior to exposure to the alarm in a separate room. For chronic CTEP dosing experiments, mice received 3 i.p. CTEP or vehicle injections, one every 48 hours, with the final dose occurring 1-2 hours prior to testing. All injections began at P23-25 (immediately following weaning). Animals were habituated to the behavioral chamber (28×17.5×12 cm transparent plastic box) for 1 minute prior to stimulus onset. The auditory stimulus was a 125 dB at 0.25 m siren (modified personal alarm, RadioShack model 49-1010, powered from a DC converter). Seizures were scored for incidence during a 2-minute stimulus presentation or until the animal reached an AGS endpoint. Wild running/jumping, status epilepticus, respiratory arrest or death were all scored as seizure activity.

### Spontaneous spiking in visual cortex

Visual cortical excitability experiments were performed as previously described [32]. 350 μm thick acute brain slices containing primary visual cortex were isolated from P20-P21 Fmr1-KO and WT littermate animals or from Fmr1-KO animals that had received chronic or acute CTEP or vehicle injections beginning at P16-P17. Slice were prepared using a Leica Vibratome in ice-cold cutting solution containing (in mM): 87 NaCl, 3 KCl, 1.25 NaH_2_PO_4_, 26 NaHCO_3_, 0.5 CaCl_2_, 7 MgCl_2_, 20 glucose, 1.3 ascorbate, 75 sucrose, saturated with 95% O_2_ and 5% CO_2_. Slices were recovered for 30 minutes at 32°C and then for an additional 2.5 hours at room temperature in a modified aCSF containing (in mM): 124 NaCl, 3.5 KCl, 1.25 NaH_2_PO_4_, 26 NaHCO_3_, 10 glucose, 0.8 MgCl_2_, and 1 CaCl_2_, saturated with 95% O_2_ and 5% CO_2_. During recordings, slices received a single 0.2 ms duration electrical stimulation of the white matter (clustered bipolar tungsten, FHC) every 30 seconds using a stimulus intensity between 40−80 μA and neuronal activity was recorded by placing a glass recording electrode (~1 MΩ resistance when filled with aCSF) in layer 5 of primary visual cortex. Extracellular recordings were first collected in vehicle conditions for 30 minutes (60 trials total), followed by 30 additional minutes in the presence of either 30 μM MPEP or 60 μM cycloheximide. All recordings were acquired with a Multiclamp 700B amplifier (Molecular Devices), amplified 1000 times, filtered between 300 Hz and 10 kHz, and digitized at 25 kHz. Spontaneous spiking events were classified as those occurring between 3.2 seconds and 30 seconds after stimulation.

### Metabolic labeling

Metabolic labeling of new protein synthesis was performed as previously described [33]. Male P28-P32 Fmr1-KO and WT littermate mice received three CTEP or vehicle i.p. injections over five days. 1-2 hours following the final injection, mice were anesthetized with isoflurane and the hippocampus was rapidly dissected into ice-cold aCSF (in mM: 124 NaCl, 3 KCl, 1.25 NaH_2_PO_4_, 26 NaHCO_3_, 10 dextrose, 1 MgCl_2_, 2 CaCl_2_, saturated with 95% O_2_ and 5% CO_2_). 500 μm thick hippocampal slices were prepared using a Stoelting Tissue Slicer and transferred into 32.5°C aCSF (saturated with 95% O_2_ and 5% CO_2_) within 5 min. Slices were incubated in aCSF undisturbed for 3 h to allow recovery of basal protein synthesis and then transferred to either aCSF containing vehicle (dH_2_O) or MTEP (1 μM), which was present for the remainder of the experiment. Actinomycin D (25 μM) was added to the chamber for 30 min to inhibit transcription after which slices were transferred to fresh aCSF containing ~10 mCi/ml [35S] Met/Cys (Perkin Elmer) for an additional 30 min. Slices were then homogenized, and labeled proteins isolated by TCA precipitation. Radiolabel incorporation was measured with a scintillation counter and samples were subjected to a protein concentration assay (Bio-Rad). Data was analyzed as counts per minute per microgram of protein, normalized to the [35S] Met/Cys aCSF used for incubation. The average incorporation of all samples was analyzed and then normalized to percent wildtype for each experiment.

### Inhibitory avoidance assay

Inhibitory avoidance (IA) experiments were performed as previously described [32]. Group housed male Fmr1-KO and WT littermate mice received three injections of 2 mg/kg CTEP or vehicle over five days beginning at ~P28. Drug was then withheld for approximately four weeks. Two days prior to IA testing, all animals were habituated to handling which consisted of scruffing mice for ~10 seconds to verify the ear tag number, followed by five minutes of resting in the gloved hands of the investigator and being allowed to freely explore while the tail was lightly restrained to prevent escape. On the day of testing, ~P60 animals were placed into the dark compartment of the IA training box (a two-chambered Perspex box consisting of a lighted safe side and a dark shock side separated by a trap door) for 30 seconds followed by 90 seconds in the light compartment for habituation. Following the habituation period, the door separating the two compartments was opened and animals were allowed to enter the dark compartment. Latency to enter following door opening was recorded (0-hour time point, collected between 8−10 a.m.); 1 animal with baseline entrance latency of greater than 120 sec. was excluded. After each animal stepped completely into the dark compartment with all four paws, the sliding door was closed and the animal received a single scrambled foot-shock (0.5mA, 2.0 sec) via electrified steel rods in the floor of the box. This foot shock intensity and duration caused each animal to vocalize and jump. Animals then remained in the dark compartment for 15−30 sec following the shock and were then placed in a fresh cage. After all members of a single cage experienced the training, mice were returned as a group to their home cage. Six to seven hours following IA training, mice received a retention test (6-hour time point, collected between 2-4 p.m.). During post-acquisition retention testing, each animal was placed in the lit compartment as in training; after a 90 sec delay, the door opened, and the latency to enter the dark compartment was recorded (cut-off time 537 sec). The order of animals run was preserved between trials. For inhibitory avoidance extinction (IAE) training, animals were allowed to explore the dark compartment of the box for 200 sec in the absence of foot-shock (animals remaining in the lit compartment after the cutoff were gently guided, using an index card, into the dark compartment); following IAE training, animals were returned to their home cages. Twenty-four hours following initial IA training, mice received a second retention test (24-hour time point, collected between 8-10 a.m.). Animals were tested in the same way as at the 6-hour time point, followed by a second 200 sec extinction trial in the dark side of the box; following training, animals were again returned to their home cages. Forty-eight hours following avoidance training, mice received a third and final retention test (48-hour time point, collected between 8-10 a.m.).

### Statistical Analysis

All experiments were performed by trained experimenters blind to genotype and drug treatment, and included same day, interleaved controls for genotype and drug treatment. All data are expressed as mean ± SEM, with n values represented in the figures and figure legends. Unless indicated otherwise, the n values stated in figures and figure legends represent numbers of animals (in experiments in which more than one measurement was taken from an animal (metabolic labeling and visual cortical spiking activity), the value representing this animal is the average of technical replicates). Differences in audiogenic seizure incidence were determined using a two-tailed Fisher’s exact test. For brain slice electrophysiology experiments, the effect of genotype or drug treatment of brain slices was determined using a paired two-tailed Student’s t test. For protein synthesis in figure 4b-c, differences between genotype and drug treatment were determined using a two-way ANOVA with Šídák’s multiple comparisons test for post-hoc analysis. For protein synthesis in figure 4d, differences between genotype and drug stimulation conditions were determined using a three-way repeated measures ANOVA and Šídák’s multiple comparisons test for post-hoc analysis. Differences between genotypes and treatment in the inhibitory avoidance assay were determined using a repeated measures two-way ANOVA with Greenhouse-Geisser correction and Tukey’s post-test for post-hoc analysis.

## Results

### Chronic mGluR5 inhibition induces treatment resistance of audiogenic seizure susceptibility which is overcome by inhibiting protein synthesis

The mGluR theory posits that glutamate acting via mGluR5 stimulates protein synthesis that is pathogenic in FXS. Early studies showed that a single dose of MPEP was sufficient to suppress audiogenic seizures in the Fmr1-KO mouse [30]. This result was later confirmed using CTEP, a NAM with increased selectivity for mGluR5 [34], as well as with several other structurally distinct mGluR5 inhibitors [35]. However, as initially suggested by Yan et al (2005), we showed in a recent study that treatment resistance develops rapidly, with as few as 3 doses of CTEP over 5 days (**Figure 1**, data reproduced from [32]).

**Figure 1.**
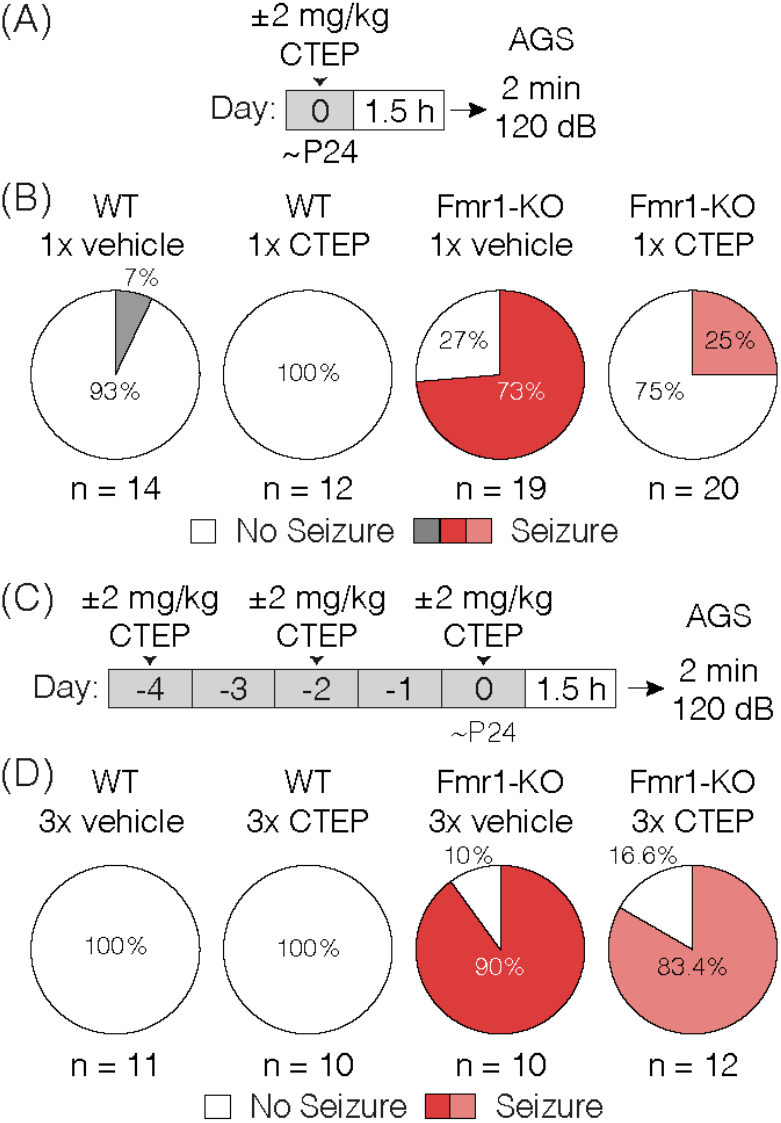
Acute but not chronic administration of the mGluR5 NAM CTEP ameliorates audiogenic seizures in Fmr1-KO mice. **(A)** Schematic shows acute CTEP dose schedule and AGS experimental design. **(B)** Fmr1-KO mice treated with vehicle exhibit increased susceptibility to audiogenic seizures compared to WT treated with vehicle (two-tailed Fisher’s exact test: p = 0.0001) and an acute dose of 2 mg/kg CTEP significantly reduced AGS incidence in Fmr1-KO mice (two-tailed Fisher’s exact test: p = 0.0038). **(C)** Schematic shows chronic CTEP dose schedule and AGS experimental design. **(D)** Chronic treatment (3 doses over 5 days) with 2 mg/kg CTEP no longer alleviates susceptibility to audiogenic seizures, indicating the development of acquired treatment resistance (Two-tailed Fisher’s exact test: Fmr1-KO vehicle treated vs. CTEP treated: p = 1.0). Data re-plotted from [32]

It is usually assumed that the effects of mGluR5 NAMs on fragile X phenotypes are due to suppression of excessive protein synthesis that occurs in the absence of FMRP. However, some fragile X phenotypes that respond to these NAMs are expressly not improved by inhibiting protein synthesis directly with cycloheximide (CHX) or other mRNA translation inhibitors [36; 37; 38; 39]. Thus, to make sense of the effects of chronic CTEP, it was important to determine if the AGS phenotype was a readout of excessive ongoing protein synthesis. To that end, we administered CHX (120 mg/kg) via a single i.p. injection in Fmr1-KO animals 1.5 hours prior to a 2-minute exposure to the 120 dB auditory stimulus. The AGS phenotype was faithfully recapitulated in vehicle-treated animals and corrected by CHX, confirming that in Fmr1-KO mice the expression of AGS susceptibility is indeed protein synthesis dependent (**Figure 2A-B**).

**Figure 2.**
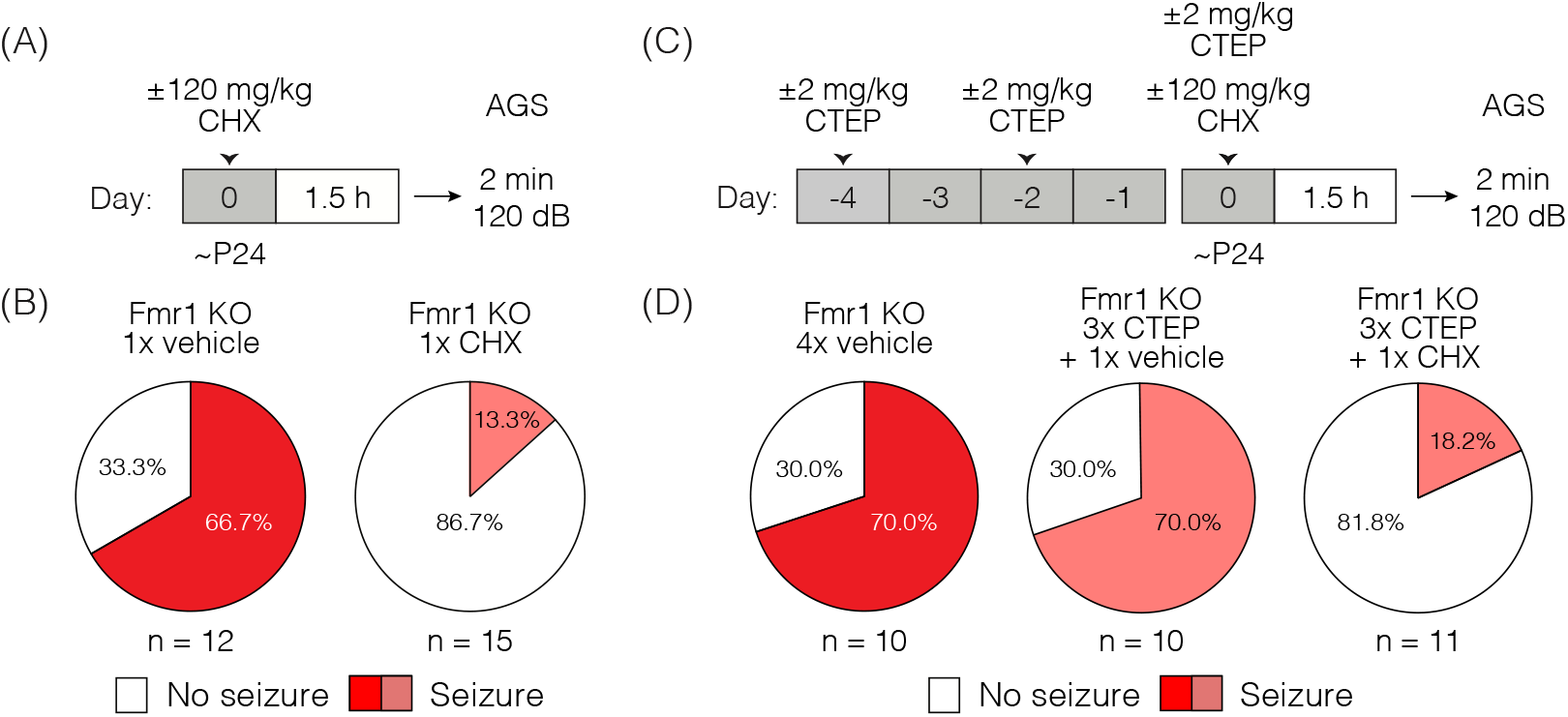
Acute cycloheximide treatment ameliorates audiogenic seizures in Fmr1-KO mice and overcomes acquired CTEP resistance. **(A)** Schematic shows acute CHX dose schedule and AGS experimental design. **(B)** Acute treatment with 120 mg/kg CHX significantly reduced AGS incidence in Fmr1-KO mice (Two-tailed Fisher’s exact test: p = 0.0069). **(C)** Schematic shows acute cycloheximide after chronic CTEP dose schedule and AGS experimental design. **(D)** Chronic CTEP (3 doses of 2 mg/kg over 5 days) causes treatment resistance that can be overcome by an acute injection of 120 mg/kg CHX immediately prior to assessing AGS (Two-tailed Fisher’s exact test: Fmr1-KO 4x vehicle vs. Fmr1-KO 3x CTEP + CHX: p = 0.0131; Fmr1-KO 3x CTEP + vehicle vs. Fmr1-KO 3x CTEP + CHX: p = 0.0131)

We next wondered if development of resistance to CTEP would also render CHX ineffective, in which case the reemergence of the AGS phenotype after chronic treatment would be explained by an entirely different pathogenic mechanism. To address this question, Fmr1-KO mice were first injected with vehicle or CTEP (2 mg/kg) 3 times over 5 days to induce treatment resistance. Then, following the third injection, mice were injected with vehicle or CHX and AGS susceptibility was measured. Fmr1-KO mice again demonstrated a robust AGS phenotype that was, as expected, not corrected by CTEP after chronic exposure (**Figure 2C-D**). However, CHX treatment was still able to acutely suppress AGS. Thus, CTEP resistance likely entails upregulation of signaling pathways that converge on protein synthesis regulation.

### Acquired treatment resistance of cortical hyperexcitability

AGS susceptibility is a complex behavioral phenotype that arises from the absence of FMRP in the inferior colliculus [40]. To explore the generality of the phenomenon of acquired treatment resistance, we employed an *in vitro* assay of neuronal hyperexcitability in visual cortical slices from Fmr1-KO mice. Previous studies have shown that layer 5 neurons display increased spontaneous spiking activity in the Fmr1-KO that is corrected acutely by treatments targeting signaling downstream of mGluR5 [12; 32]. We confirmed this cellular phenotype (**Figure 3A-B**) and then investigated the sensitivity to acute exposure to the mGluR5 NAM MPEP in Fmr1-KO mice following treatment *in vivo* with either vehicle or 3x CTEP. In brain slices prepared from Fmr1-KO animals injected with vehicle, bath application of the mGluR5 inhibitor MPEP (30 μM) rapidly (within 30 minutes) reduced the number of action potentials (**Figure 3C-E**). If animals were first treated *in vivo* with a single dose of CTEP shortly before brain slice preparation, we also observed a complete suppression of aberrant spiking activity. Inhibition of mGluR5 by CTEP is known to be long-lasting and survive slice preparation, even a day later [34]. Therefore, it was not surprising that we observed no additional suppression of spiking by bath applied MPEP in these experiments, presumably because mGluR5 was fully inhibited by the CTEP treatment (**Figure 3D-E**).

**Figure 3.**
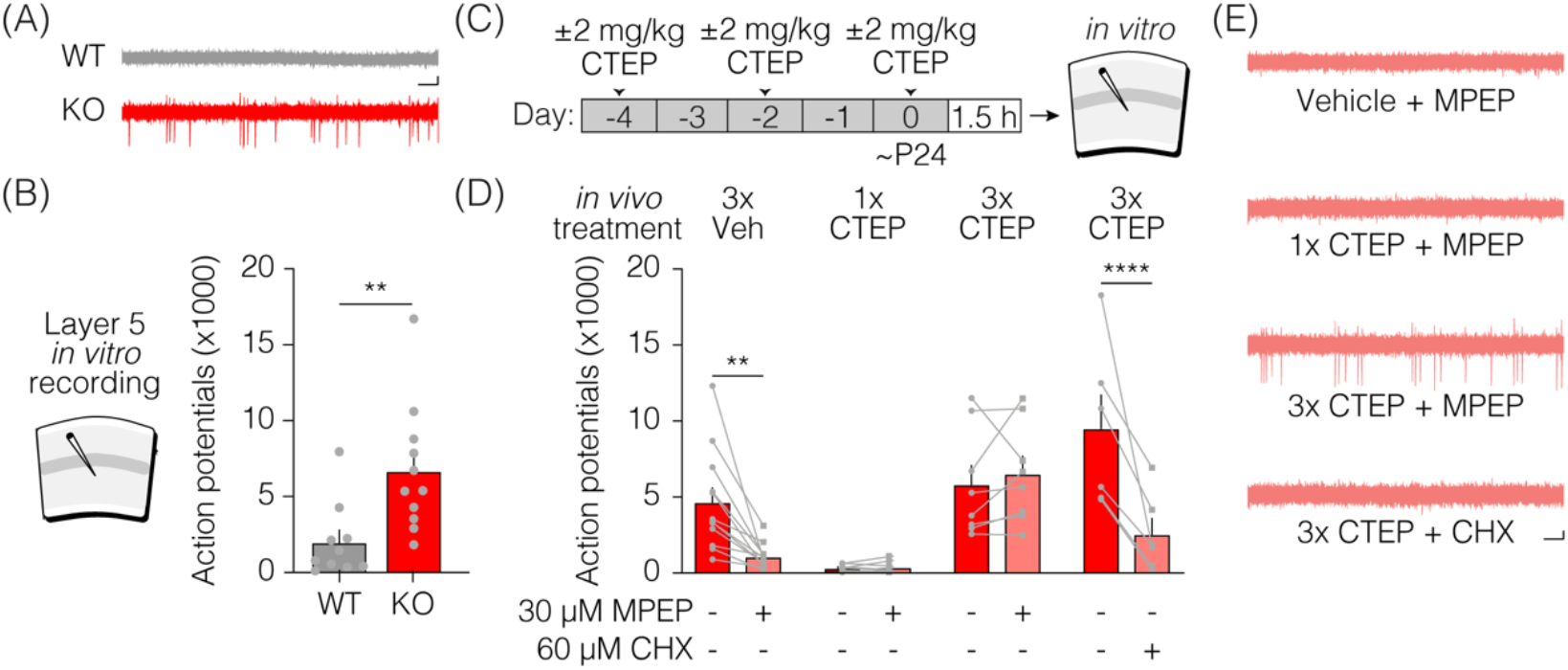
Acute but not chronic CTEP treatment ameliorates hyperexcitability in Fmr1-KO layer 5 of primary visual cortex. **(A)** Representative traces of extracellular recordings in layer 5 of primary visual cortex from wild type (black) and Fmr1-KO (red) animals show increased spontaneous firing in Fmr1-KO slices (Scale bar represents 200 μV by 200 ms). **(B)** Layer 5 neurons in Fmr1-KO mice have significantly increased spontaneous action potentials compared to wild type littermates (paired t-test: wild type vs. Fmr1-KO: p = 0.0062) **(C)** Schematic shows acute and chronic CTEP dose schedules and visual cortical excitability experimental design. **(D1)** Elevated spontaneous activity in layer 5 primary visual cortical slices from Fmr1-KO mice is significantly reduced by bath application of 30 μM MPEP (Two-tailed paired t-test: Fmr1-KO 3x vehicle + vehicle vs. Fmr1-KO 3x vehicle + MPEP: p = 0.0024) or **(D2)** an acute injection in vivo of 2 mg/kg CTEP (Two-tailed paired t-test: Fmr1-KO 1x CTEP + vehicle vs. Fmr1-KO 1x CTEP + MPEP: p = 0.1693). **(D3)** Chronic 2mg/kg CTEP (3 doses over 5 days) leads to treatment resistance of spontaneous activity in layer 5 primary visual cortex that is not overcome by bath application of 30 μM MPEP (Two-tailed paired t-test: Fmr1-KO 3x CTEP + vehicle vs. Fmr1-KO 3x CTEP + MPEP: p = 0.4327) but **(D4)** is significantly reduced by bath application of 60 μM CHX (Two-tailed paired t-test: Fmr1-KO 3x CTEP + vehicle vs. Fmr1-KO 3x CTEP + CHX: p = 0.0102). **(E)** Representative traces of extracellular recordings in layer 5 primary visual cortex showing spontaneous activity in Fmr1-KO slices treated with bath applied 30 μM MPEP, animals injected acutely with 2mg/kg CTEP, and animals injected chronically (3 doses over 5 days) with 2 mg/kg CTEP followed by bath application of either 30 μM MPEP or 60 μM CHX (Scale bar represents 200 μV by 200 ms). Data are displayed as mean ±SEM.

The findings were very different after 3 doses of CTEP *in vivo*. Layer 5 neurons displayed the characteristic hyperexcitability phenotype which could no longer be reversed by MPEP. The inability of bath applied MPEP (on top of the residual CTEP in the tissue) to correct the hyperexcitability suggests that the molecular mechanism of acquired treatment resistance is unlikely to be explained simply by an upregulation of receptors in the membrane. However, as was observed for the AGS phenotype, inhibition of protein synthesis with CHX was still effective in correcting this phenotype even after the development of resistance to the mGluR5 NAMs (**Figure 3D-E**).

### Chronic mGluR5 inhibition fails to normalize elevated hippocampal protein synthesis rates in Fmr1-KO mice

An elevated rate of basal protein synthesis in the hippocampus of Fmr1-KO mice and rats is a hallmark fragile X phenotype, and it is rescued by inhibiting proteins in the signaling cascade linking mGluR5 activation with protein synthesis [5; 12; 32; 33; 41; 42; 43; 44; 45]. Consistent with these prior studies, we found that a single dose of CTEP *in vivo* corrects the protein synthesis phenotype measured ~4 h later in hippocampal slices (**Figure 4A-B**). Once again, this effect was lost after dosing CTEP 3 times over 5 days (**Figure 4C**). Thus, the acquired resistance to treatment with the mGluR5 NAM was not restricted to excitability phenotypes, but also generalizes to a core biochemical phenotype in another brain region.

**Figure 4.**
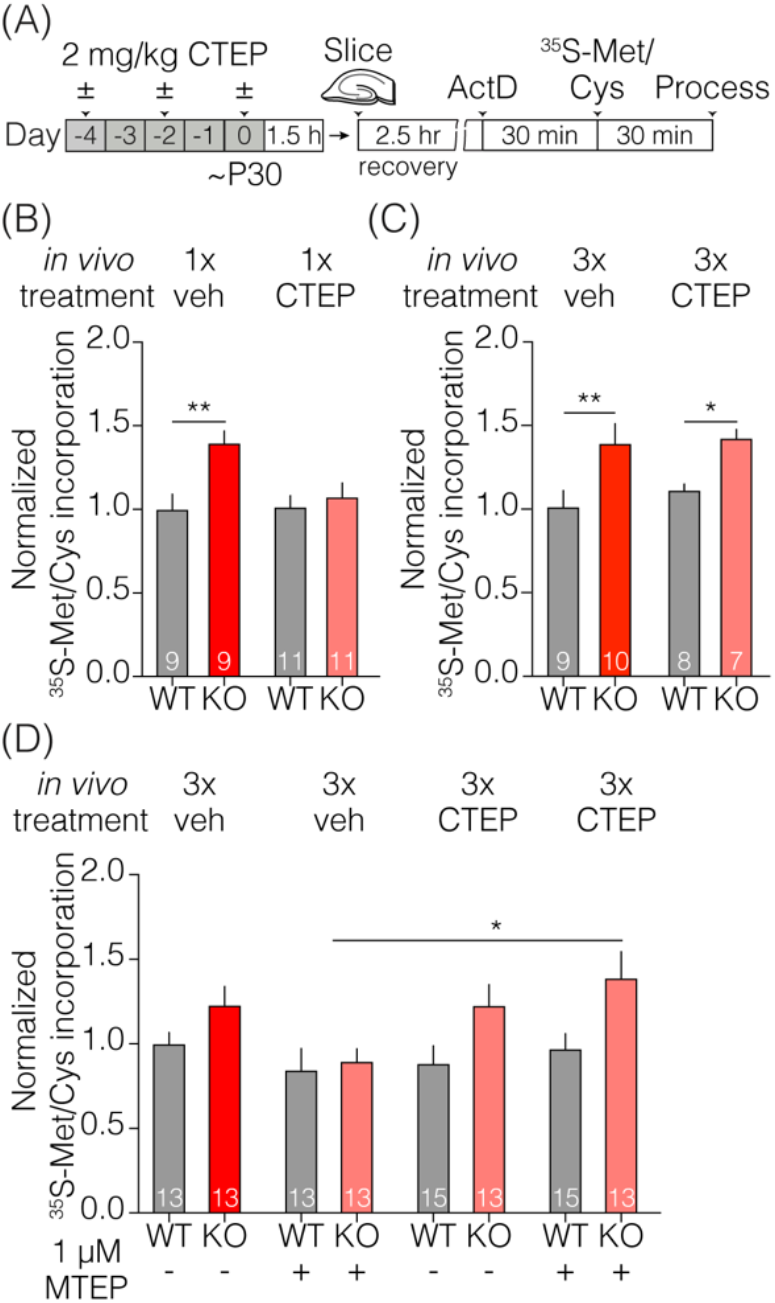
Acute but not chronic CTEP treatment ameliorates elevated basal protein synthesis rates in Fmr1-KO hippocampal slices. **(A)** Schematic shows acute and chronic CTEP dose schedules and hippocampal metabolic labeling experimental design. **(B)** Basal protein synthesis rates are increased in hippocampal slices prepared from Fmr1-KO mice compared to wildtype littermate animals and acute *in vivo* treatment with 2 mg/kg CTEP restores Fmr1-KO protein synthesis rates to wildtype levels. There was a statistically significant effect of genotype (two-way ANOVA, F (1, 36) = 8.341, p = 0.0066) and a significant interaction between genotype and treatment. (two-way ANOVA, F (1, 36) = 4.501, p = 0.0408; Šídák’s multiple comparisons test: wild type 1x vehicle vs. Fmr1-KO 1x vehicle p = 0.0036, wild type 1x CTEP vs. Fmr1-KO 1x CTEP p = 0.8183). **(C)** Chronic (3 doses over 5 days) *in vivo* treatment with 2 mg/kg CTEP has no effect on radiolabel incorporation in hippocampal slices from wildtype or Fmr1-KO animals. There was a statistically significant effect of genotype (two-way ANOVA, F (1, 30) = 15.13, p = 0.0005; Šídák’s multiple comparisons test: wild type 3x vehicle vs. Fmr1-KO 3x vehicle p = 0.0082, wild type 3x CTEP vs. Fmr1-KO 3x CTEP p = 0.0410). **(D)** Bath application of 1 μM MTEP reduces elevated basal protein synthesis rates in Fmr1-KO hippocampal slices to wild type levels but has no effect on hippocampal slices from animals injected with chronic (3 doses over 5 days) 2.0 mg/kg CTEP. There was a statistically significant effect of genotype (three-way ANOVA, F (1, 50) = 8.013, p = 0.0067) and a significant interaction between *in vivo* treatment and *in vitro* treatment. (three-way ANOVA, F (1, 50) = 8.536, p = 0.0052; Šídák’s multiple comparisons test: Fmr1-KO 3x vehicle + MTEP vs. Fmr1-KO 3x CTEP + MTEP p = 0.0347). Data are displayed as mean ±SEM.

We next asked if supplemental bath application of a second mGluR5 NAM could overcome the resistance observed after repeated dosing with CTEP treatment *in vivo*. In these experiments we used MTEP, which is more selective and potent than MPEP [46; 47]. Although MTEP reversed the fragile X phenotype in animals treated previously with vehicle, this effect was lost in animals treated chronically with CTEP (**Figure 4D**). This finding again suggests that this phenomenon is not due to changes in the expression of the mGluR5 receptor but rather depends on an adaption of intracellular signaling.

### The mechanism of acquired treatment resistance lies downstream of GSK3α activation

The signaling pathways that link mGluR5 activation with the regulation of protein synthesis are well described [27]. The Ras-ERK1/2 pathway is of considerable interest in FXS, as inhibitors such as lovastatin and metformin that target this signaling arm correct myriad phenotypes in the Fmr1-KO mouse and rat (**Figure 5A**) [12; 48; 49], as well as biochemical phenotypes in platelets and neurons derived from patients with FXS [43; 50]. We therefore could elucidate where in this signaling cascade acquired treatment resistance arises by testing which interventions more proximal to protein synthesis can overcome the effects of chronic CTEP. We recently demonstrated that the paralog-specific GSK3α inhibitor BRD0705 corrects AGS susceptibility, cortical hyperexcitability, and basal protein synthesis phenotypes in Fmr1-KO animals [32]. In wildtype mice, BRD0705 inhibits the stimulation of protein synthesis by mGluR5 activation, but not phosphorylation of ERK1/2, thus indicating that it acts well downstream in the signaling pathway. We therefore reasoned that by acting more proximal to protein synthesis regulation, BRD0705 might be able to overcome the acquired treatment resistance induced by chronic CTEP.

**Figure 5.**
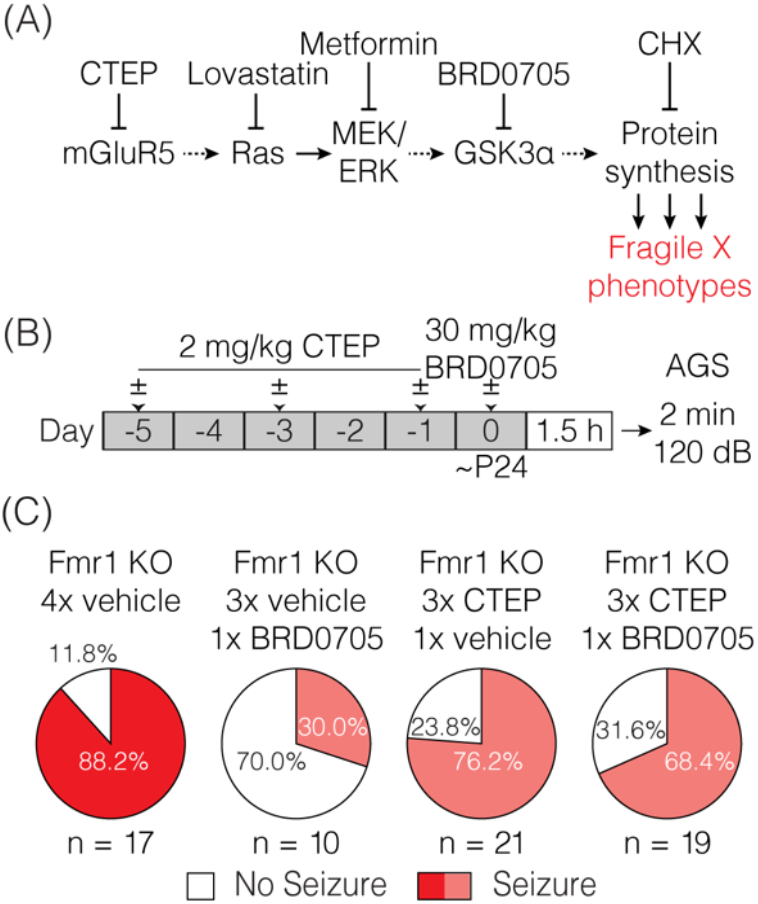
Inhibiting GSK3α does not overcome audiogenic seizure treatment resistance induced by chronic CTEP. **(A)** Some elements of the signaling pathway that couples mGluR5 to protein synthesis. AGS susceptibility in fragile X can be corrected by the protein synthesis inhibitor CHX and by compounds acting at several different nodes in this pathway, including GSK3α. **(B)** Schematic shows drug dosing schedule and AGS experimental design. **(C)** Chronic (3 doses over 5 days) treatment with 2.0 mg/kg CTEP followed by a vehicle injection does not alter Fmr1-KO audiogenic seizure susceptibility (two-tailed Fisher’s exact test: Fmr1-KO 4x vehicle vs. Fmr1-KO 3x CTEP +1x vehicle p = 0.4267). A single injection of 30 mg/kg BRD0705 normalizes audiogenic seizure susceptibility in Fmr1-KO mice but has no effect on seizure incidence in Fmr1-KO mice treated with chronic (3 doses over 5 days) 2.0 mg/kg CTEP (two-tailed Fisher’s exact test: Fmr1-KO 4x vehicle vs. Fmr1-KO 3x vehicle + BRD0705 p = 0.0036; two-tailed Fisher’s exact test: Fmr1-KO 4x vehicle vs. Fmr1-KO 3x CTEP + BRD0705 p = 0.2357).

To test this hypothesis, resistance to treatment with mGluR5 NAMs was first induced by 3x CTEP injections and then the effect of BRD0705 was examined. In controls, receiving only vehicle before BRD0705, we confirmed the suppression of AGS by inhibiting GSK3α. However, previous chronic exposure to CTEP eliminated the ameliorative effect of BRD0705 (**Figure 5B-C**). These findings suggest that whatever cellular adaption is responsible for acquisition of treatment resistance following CTEP treatment, it is likely to occur downstream of signaling by GSK3α (but upstream of protein synthesis; see **Figure 2D**).

### Temporary early intervention with CTEP corrects inhibitory avoidance behavior one month after treatment

We have demonstrated the development of rapid treatment resistance in audiogenic seizure susceptibility, visual cortical hyperexcitability, and hippocampal protein synthesis, after only 3 doses of CTEP. However, a previous study showed that a deficit in inhibitory avoidance (IA), a contextual learning behavior, was corrected in mice receiving doses of CTEP (2 mg/kg p.o.) every other day for >1 month (achieving a steady state receptor occupancy of >80%)[34]. Clearly that chronic dosing schedule would have caused treatment resistance in the assays employed here.

How might we reconcile these disparate findings? A recent study using the rat model of fragile X offers a clue [48]. They demonstrated that temporary administration of lovastatin for 5 weeks starting at one month of age produced a persistent improvement of cognitive behavior that could be measured many weeks after the drug was discontinued. These findings suggest that brief interventions during a postnatal critical period may be sufficient to produce lasting effects. Thus, we wondered if the effect of CTEP on IA memory observed previously by Michalon et al. (2012) was actually not a consequence of continuous inhibition of mGluR5, but rather was due to the fortuitous timing of the first dose(s) of CTEP.

To assess this hypothesis, we treated Fmr1-KO mice and wild type littermates with vehicle or CTEP (2 mg/kg i.p.) three times over five days beginning at postnatal day 28 and then withheld the drug for four weeks and measured inhibitory avoidance behavior beginning at postnatal day 58 (**Figure 6A**). Vehicle injected Fmr1-KO animals replicated the well described IA deficits in memory acquisition and extinction compared to vehicle treated wild type littermates (**Figure 6B**). However, early and temporary intervention with CTEP fully restored memory acquisition and extinction in Fmr1-KO mice, providing evidence for a critical window in which mGluR5 inhibition produces a durable behavioral improvement. These findings suggest that the long-term treatment regimen deployed in previous studies was unnecessary for the behavioral rescue.

**Figure 6.**
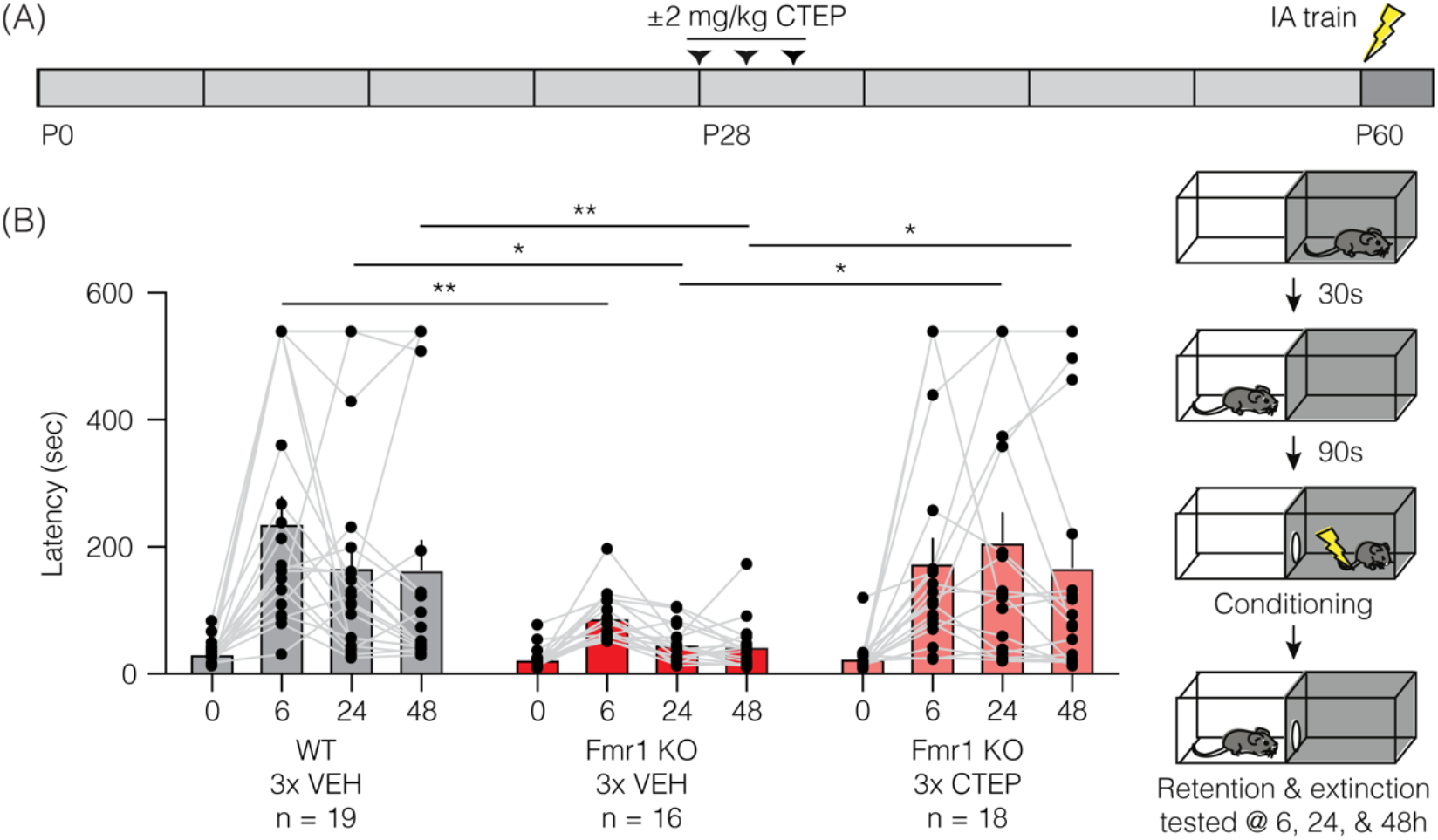
Brief treatment of juvenile Fmr1-KO mice with CTEP normalizes inhibitory avoidance measured one month after the end of treatment. **(A)** Schematic shows when during the developmental timeline CTEP is administered and IA experimental design. **(B)** There was a statistically significant interaction between treatment and time point (repeated measures two-way ANOVA with Greenhouse-Geisser correction, F (6, 150) = 2.684, p = 0.0064). Fmr1-KO mice treated with vehicle displayed impaired acquisition and extinction of IA learning compared to vehicle treated wild type mice (wild type vehicle vs. Fmr1-KO vehicle, Tukey’s post-test at time 0 hour p = 0.3488, at 6 hours p = 0.0048, at 24 hours p = 0.0116, at 48 hours p = 0.0444) and CTEP treated Fmr1-KO mice (Fmr1-KO vehicle vs. Fmr1-KO 3x CTEP, Tukey’s post-test at time 0 hour p = 0.9756, at 6 hours p = 0.1024, at 24 hours p = 0.0067, at 48 hours p = 0.0416). Data are displayed as mean ±SEM.

## Discussion

Our results confirm the previous observation of acquired resistance to mGluR5 inhibition in the AGS assay [30; 31; 32] and demonstrate that this applies to two additional fragile X phenotypes: cortical hyperexcitability and exaggerated protein synthesis. These behavioral, cellular, and biochemical phenotypes arise from dysfunction in three different parts of the brain— inferior colliculus, primary visual cortex, and hippocampus—suggesting that acquired treatment resistance is a general phenomenon, not restricted to one part of the nervous system or one functional readout.

One feature that these fragile X phenotypes have in common, however, is that they have been shown to be corrected rapidly (within tens of minutes) by acute inhibition of mGluR5 signaling in drug naïve animals [12; 30; 33]. This rapid drug response has historically been attributed to suppression of mGluR5-regulated protein synthesis. In support of this interpretation, we show here (to our knowledge, for the first time) that both increased susceptibility to AGS and cortical hyperexcitability are also reversed acutely by CHX, an inhibitor of mRNA translation (**Figures 2, 3**). These findings suggest the existence of pathogenic protein species that are rapidly depleted by inhibiting mGluR5 or protein synthesis. Identifying these protein(s) will be of great interest as they represent potential therapeutic targets that might allow for more precise molecular interventions that do not require manipulating proteostasis directly.

At this point, the mechanism for the acquired treatment resistance is unknown, but our experiments help to narrow the possibilities. We note that use of the term “tolerance” has been avoided in this paper, because it is typically used to describe the reduced effectiveness of receptor *agonists* with prolonged exposure. Desensitization of G-protein coupled receptors upon ligand binding is a well-known phenomenon, and is accounted for by changes in receptor surface expression and/or the coupling of the receptors to their G-proteins [51; 52; 53]. It is possible that an inverse process—sensitization or increased expression of mGluR5—after chronic NAM treatment accounts for the diminished effectiveness of treatment, but several lines of evidence suggest otherwise. First, in the visual cortex hyperexcitability and hippocampal protein synthesis assays, the lost effectiveness of acute CTEP after chronic dosing was not mitigated by the bath application of another mGluR5 NAM (**Figure 3, 4**). Considered with the previous finding that mGluR5 protein expression is not increased after development of NAM resistance in the AGS assay [31], these data suggest that the mechanism lies downstream of the receptor. This interpretation is supported by the additional finding that resistance to CTEP blocks the effectiveness of inhibiting GSK3α, an enzyme believed to act downstream of ERK in the pathway coupling mGluR5 to protein synthesis (**Figure 5**)[32]. Thus, the treatment resistance we observe here is best conceptualized as an intracellular adaptation to chronic loss of signaling through mGluR5. The re-expression of pathogenic proteins that cause neuronal hyperexcitability after chronic CTEP presumably is the ultimate basis for acquired treatment resistance, because phenotypic rescue is still possible using a protein synthesis inhibitor.

A previous study did not assess the effects of chronic CTEP administration on hippocampal protein synthesis but clearly demonstrated that chronic, weeks long, CTEP dosing fully restored many other phenotypes including inhibitory avoidance, dendritic spine morphology, and overactivity of ERK and mTOR [34]. One important clue to resolving this discrepancy comes from our finding that early and temporary modulation of mGluR5 with CTEP rescues inhibitory avoidance behavior measured four weeks after withholding CTEP. There now exists an emerging body of evidence that there may be critical developmental windows during which manipulating mGluR5 signaling may restore normal developmental trajectories leading to long lasting behavioral improvements. A recent series of experiments using the Ras/ERK inhibitor lovastatin showed that temporary treatment early in juvenile development of Fmr1-KO rats corrected learning and memory behavioral deficits weeks after the drug had been withheld [48].

Taken together, these data lead to the hypothesis that the sustained correction of some phenotypes observed after chronic CTEP may have been due to the restoration of a normal developmental trajectory early in treatment, rather than reflecting the need for sustained modulation of mGluR5 signaling throughout postnatal maturation (**Figure 7A**). Future studies must continue to refine the timing of these early therapeutic windows and determine if they are specific to individual phenotypes or if they permit the correction of a wide range of fragile X phenotypes. More broadly, these new findings highlight the utility of model systems like the Fmr1-KO mouse and rat to fully examine how the timing of initiation and duration of treatments affects behavioral rescue outcomes.

**Figure 7.**
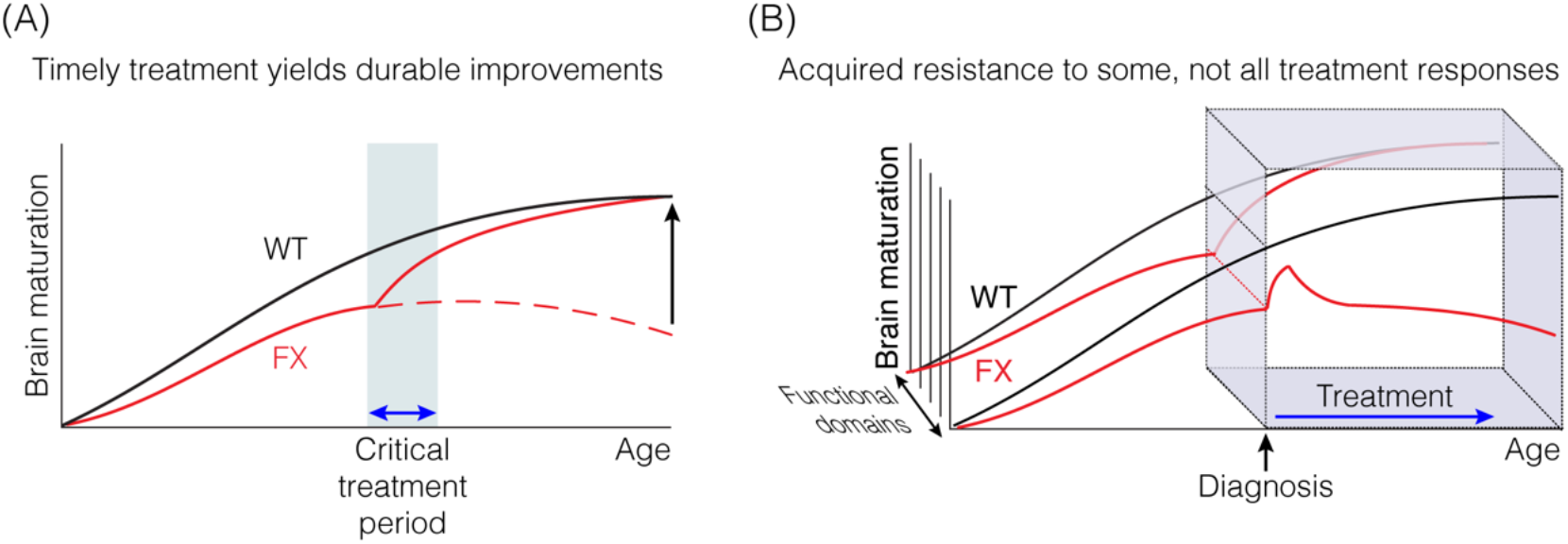
Timely therapeutic interventions have the potential to correct the derailed postnatal development of some cognitive functions. **(A)** Some improvements may not be immediately apparent, but emerge with time after treatment is discontinued. **(B)** Some measures of improvement may not be as susceptible to acquired treatment resistance as others.

Recognition of both the risk of acquired treatment resistance, and the opportunity of inducing a persistent improvement with timely intervention during development, should inform the design of future clinical trials. These insights are also usefully applied to the interpretation of the previous trials in FXS using mGluR5 NAMs. The first thing to note is that acquired treatment resistance is not an inevitable consequence of inhibiting mGluR5 signaling. For example, although we saw profound loss of effectiveness in the AGS assay after only 3 doses of CTEP over 5 days, BRD0705 and MPEP retain anticonvulsant activity after 5-6 daily doses [31; 32]. Both compounds have a far shorter half-life than CTEP, and daily dosing does not produce continuous inhibition of mGluR5 signaling. Thus, pulsatile rather than sustained inhibition of mGluR5 might have durable therapeutic efficacy while avoiding maladaptive treatment resistance. We note that there are considerable differences in the pharmacokinetic properties of mGluR5 NAMs that have advanced to human studies, with affinities that range over 200-fold [54]. In published FXS trials, mavoglurant (AFQ056), with a half-life of ~12 hours [55], was dosed twice daily for 12 weeks [26]. It is entirely possible that the failure of these trials to demonstrate benefit in their primary endpoint was due to acquired treatment resistance. It is also possible that not all efficacy measures are equally affected by treatment resistance (**Figure 7B**), perhaps explaining why significant improvements could still be observed using other functional measures, notably in subjects receiving the lower drug doses [56]. Regardless, our findings make the case for limiting the duration of continuous treatment, perhaps by providing rest periods, to prevent development of drug resistance that may have masked some benefits of mGluR5 NAMs in previous clinical trials.

Another factor that clearly needs to be considered is the age at treatment onset. Our findings together with those in the rat model of fragile X [48] suggest that timely postnatal intervention may be sufficient to restore some cognitive functions by correcting the trajectory of subsequent brain circuit development. The youngest age in the mavoglurant clinical trial was 12 years old. It is possible that this age of treatment onset was too late, or that the corrective effect of treatment on brain development takes longer than 12 weeks to be detectable, as has been suggested by observations during the open-label extension period lasting over 2 years [57]. Taken together, the evidence indicates that it is premature to abandon the mGluR theory as an organizing principle for developing new disease-modifying therapies in FXS.

## Conflict of Interest

Mark Bear is a founder of Allos Pharma, developing treatments for fragile X

## Author Contributions

PKM, DCS, RKS, AJH, and MFB conceived of the experiments. PKM conducted and analysed the hippocampal protein synthesis experiments. PKM, RKS, and AJH conducted and analysed the AGS experiments. DCS conducted and analysed the inhibitory avoidance experiment. RKS conducted and analysed the extracellular recording experiments. DCS, AJH, and MFB prepared the manuscript.

## Funding

This work was supported by R01MH106469 (MFB), the Picower Institute Innovation Fund (MFB), FRAXA postdoctoral fellowships (RKS and PKM), and T32MH112510 (DCS).

## Acknowledgments

We thank Amanda Coronado, Erin Hickey, and Kiki Chu for excellent technical assistance and laboratory support. We are also pleased to acknowledge Jessica Buckey and Athene Wilson-Glover for administrative assistance.

## Data Availability Statement

All data associated with this paper are in the main text or supplementary materials.

